# Gamma activity accelerates during prefrontal development

**DOI:** 10.1101/2020.03.10.986281

**Authors:** Sebastian H. Bitzenhofer, Jastyn A. Pöpplau, Ileana L. Hanganu-Opatz

## Abstract

Gamma oscillations are a prominent activity pattern in the cerebral cortex. While gamma rhythms have been extensively studied in the adult prefrontal cortex in the context of cognitive (dys)functions, little is known about their development. We addressed this issue by using extracellular recordings and optogenetic stimulations in mice across postnatal development. We show that fast rhythmic activity in the prefrontal cortex becomes prominent during the second postnatal week. While initially at about 15 Hz, fast oscillatory activity progressively accelerates with age and stabilizes within gamma frequency range (30-80 Hz) during the fourth postnatal week. Activation of layer 2/3 pyramidal neurons drives fast oscillations throughout development, yet the acceleration of their frequency follows similar temporal dynamics as the maturation of fast-spiking interneurons. These findings uncover the development of prefrontal gamma activity and provide a framework to examine the origin of abnormal gamma activity in neurodevelopmental disorders.

## Introduction

Synchronization of neuronal activity in fast oscillatory rhythms is a commonly observed feature in the adult cerebral cortex. While its exact functions are still a matter of debate, oscillatory activity in gamma frequency range has been proposed to organize neuronal ensembles and to shape information processing in cortical networks^1–3^. Gamma activity emerges from reciprocal interactions between excitatory and inhibitory neurons. Fast inhibitory feedback via soma-targeting parvalbumin (PV)-expressing inhibitory interneurons leads to fast gamma activity (30-80 Hz)^4,5^, whereas dendrite-targeting somatostatin (SOM)-expressing inhibitory interneurons contribute to beta/low gamma activity (20-40 Hz)^5,6^. A fine-tuned balance between excitatory drive and inhibitory feedback is mandatory for circuit function underlying cognitive performance. Interneuronal dysfunction and ensuing abnormal gamma activity in the medial prefrontal cortex (mPFC) have been linked to impaired cognitive flexibility^7^. Moreover, imbalance between excitation and inhibition in cortical networks and resulting gamma disruption have been proposed to cause cognitive disabilities in autism and schizophrenia^7–9^.

Despite substantial literature linking cognitive abilities and disabilities to gamma oscillations in the adult mPFC, the ontogeny of prefrontal gamma activity is poorly understood. This knowledge gap is even more striking considering that abnormal patterns of fast oscillatory activity have been described at early postnatal age in autism and schizophrenia mouse models^10–12^. Knowing the time course of prefrontal gamma maturation is essential for understanding the developmental aspects of mental disorders.

To this end, we performed an in-depth investigation of the developmental profile of gamma activity in the mouse mPFC from postnatal day (P) 5 until P40. We show that pronounced fast oscillatory activity emerges towards the end of the second postnatal week and increases in frequency and amplitude with age. While activation of layer 2/3 pyramidal neurons (L2/3 PYRs) drives fast oscillatory activity throughout development, the acceleration of its frequency follows the same dynamics as the maturation of inhibitory feedback and fast-spiking interneurons.

## Results

### Fast oscillatory activity in the prefrontal cortex accelerates during development

Extracellular recordings in the mPFC of anesthetized and non-anesthetized P5-40 mice revealed that oscillatory activity at fast frequencies (>12 Hz) can be detected at the beginning of the second postnatal week. The temporal dynamics of these fast oscillations is similar in the two states, yet their magnitude is higher in awake mice, as described in previous studies^13^. The magnitude of fast oscillations increases with age (Mann-Kendall trend test, p=3.93*10^−22^, n=114 recordings, tau-b 0.625) and can be detected as distinct peaks in power spectra at the end of the second postnatal week (Figure 1a,b). The peak frequency of these oscillations gradually increases with age (Mann-Kendall trend test, p=2.73*10^−8^, n=114 recordings, tau-b 0.361), starting at ~20 Hz at P12 and reaching the characteristic gamma frequency of 50-60 Hz at P25 (Figure 1b-d). Both, peak strength and peak frequency, do not change after P25.

**Figure 1.**
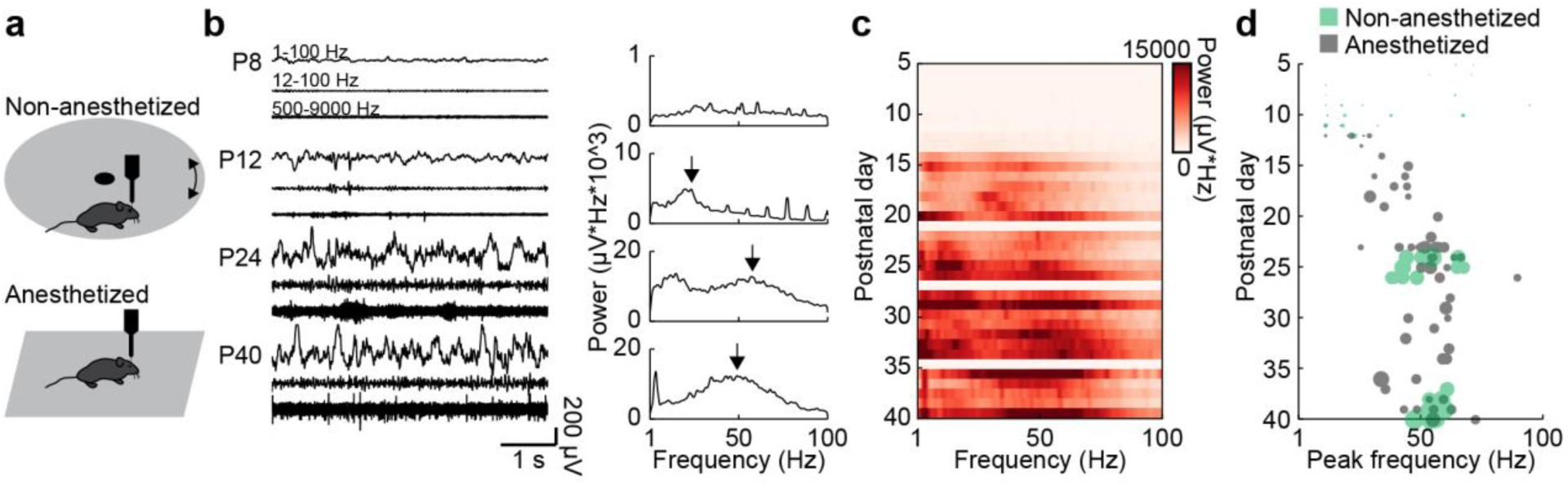
Development of gamma activity in the mouse mPFC. (**a**) Schematic of extracellular recordings in the mPFC of anesthetized and non-anesthetized P5-40 mice. (**b**) Characteristic extracellular recordings of local field potentials (LFP) and multi-unit activity (MUA) at different ages after band-pass filtering (left) and the corresponding power spectra (right). (**c**) Color-coded average power spectra of spontaneous oscillatory activity for P5-40 mice. (**d**) Scatter plot displaying peak frequencies of fast oscillations (12-100 Hz) during postnatal development of anesthetized (gray, n=80) and non-anesthetized mice (green, n=20, 35 recordings). Marker size displayed peak strength. (See supplementary table 1 for statistics.)

### Fast spiking interneuron maturation resembles the time course of gamma development

Fast spiking (FS) PV-expressing interneurons have been identified as key elements for the generation of oscillatory activity in gamma frequency range in the adult cortex^4^. To assess whether the developmental gamma dynamics relate to FS PV-expressing interneuron maturation, we performed immunohistochemistry and single unit analysis in P5-40 mice.

First, immunostainings showed that PV expression in the mPFC starts at the end of the second postnatal week, increases until P25 and stabilizes afterwards (Mann-Kendall trend test, p=1.29*10^−7^, n=38 slices, tau-b 0.623) (Figure 2a). This dynamics over age mirrors the changes in peak power and peak frequency of the fast oscillations described above. In contrast, the number of SOM positive neurons does not significantly vary along postnatal development (Mann-Kendall trend test, p=0.99, n=39 slices, tau-b - 0.003) (Figure 2b).

**Figure 2.**
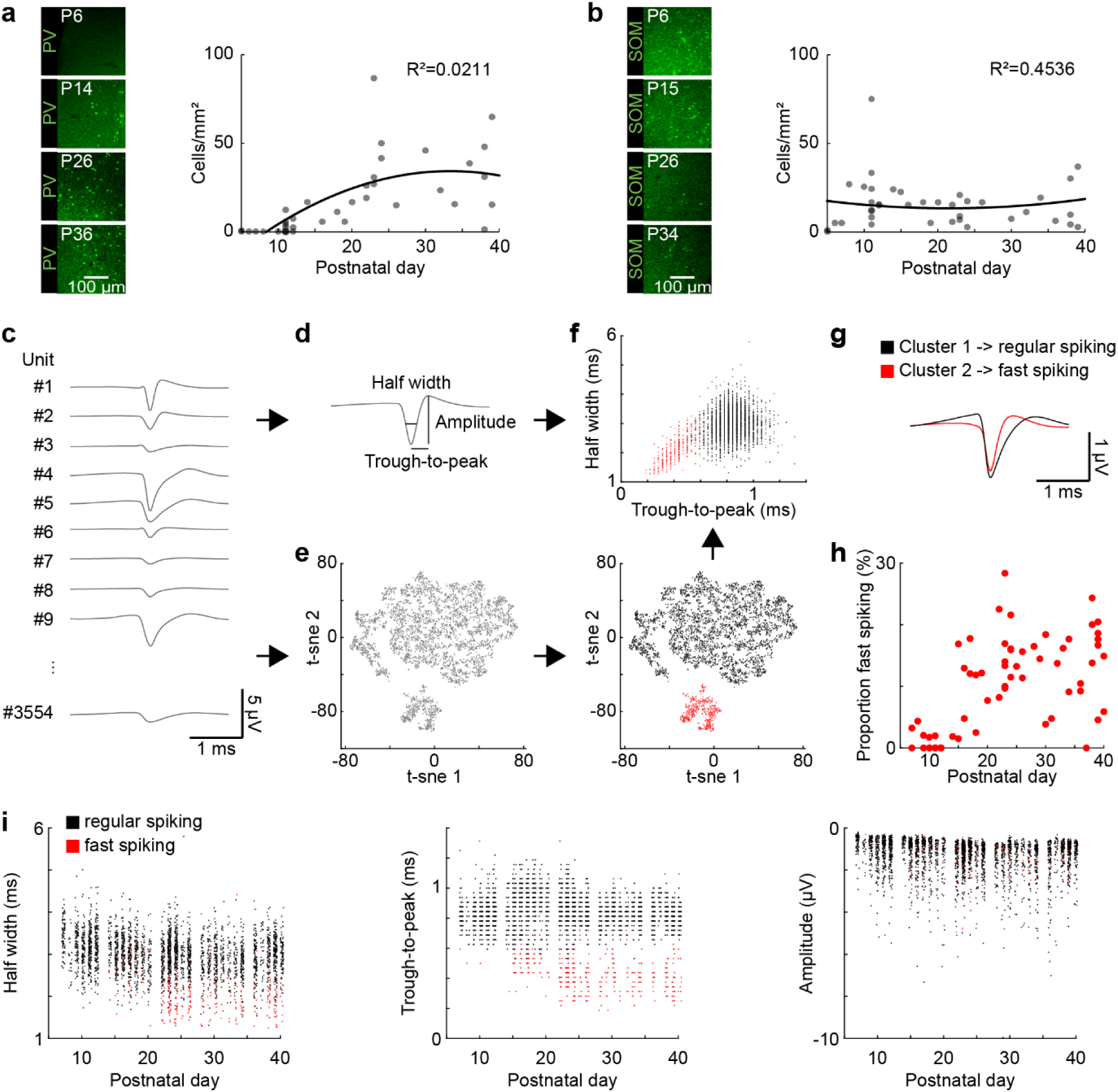
Development of FS interneurons in the mouse mPFC. (**a**) Left, examples of PV immunostaining in the mPFC at different ages. Right, scatter plot displaying the density of PV-immunopositive neurons in the mPFC of P5-40 mice (n=38). (**b**) Same as (a) for SOM-immunopositive neurons (n=39). (**c**) Example mean waveforms of extracellular recorded units from P5-40 mice. (**d**) Schematic showing features classically used to distinguish RS and FS units in adult mice. (**e**) Left, scatter plot showing the first two components of a t-sne dimensionality reduction on the mean waveforms for all units recorded from P5-40 mice. Right, same as left with the first two clusters obtained by hierarchical clustering labeled in black and red. (**f**) Scatter plot of half width and trough to peak time cluster 1 (black) and 2 (red). (**g**) Mean waveform for cluster 1 (black) and 2 (red). (**h**) Scatter plot showing the proportion of FS units for P5-40 mice. (**i**) Scatter plots showing classic spike shape features for P5-40 for cluster 1 (RS, black) and 2 (FS, red). (Average data is presented as median ± MAD. See supplementary table 1 for statistics.)

Second, to directly assess the functional maturation of FS PV-expressing neurons, we used the extracellular recordings of multi-unit activity and identified single units. The classically used action potential features to distinguish adult FS and regular spiking (RS) neurons (i.e. trough to peak time and half width) cannot be applied during early development, because of a strong overlap of these features. Therefore, we developed an algorithm to classify RS and FS using dimensionality reduction with t-Distributed Stochastic Neighbor Embedding (t-sne) on the mean waveforms of all units recorded across development, followed by hierarchical clustering (Figure 2c-e). This approach results in an unbiased detection of FS units across age (Figure 2f,g). Of note, a more detailed analysis reveals that the dimensionality reduction based on mean waveforms correlates with features like trough to peak and half width, but is less affected by age, layer and amplitude (Figure 2 – figure supplement 1). The classification of units with this method reveals that FS units start to be detected at the end of the second postnatal week. Their number gradually increased until P25 (Mann-Kendall trend test, p=1.44*10^−7^, n=66 recordings, tau-b 0.458) (Figure 2h). A comparison of the classical features across age showed that trough-to-peak duration (Mann-Kendall trend test, RS, p=4.57*10^−5^, n=3172 units, tau-b -0.051; FS, p=9.23*10^−11^, n=382 units, tau-b -0.236), half width (Mann-Kendall trend test, RS, p=1.61*10^−28^, n=3172 units, tau-b -0.134; FS, p=5.17*10^−17^, n=382 units, tau-b -0.295), and negative amplitude (Mann-Kendall trend test, RS, p=4.45*10^−22^, n=3172 units, tau-b -0.117; FS, p=3.82*10^−6^, n=382 units, tau-b -0.163) of RS and FS units gradually decreased from the end of the second postnatal week until P25. However, the most prominent changes were detected for through-to-peak duration and half width of FS units (Figure 2i).

Thus, in line with the immunohistochemical examination, the analysis of single units showed that FS putatively PV-positive interneurons show similar dynamics of maturation as fast oscillations recorded in the mPFC of P5-40 mice.

**Figure 2 – figure supplement 1.**
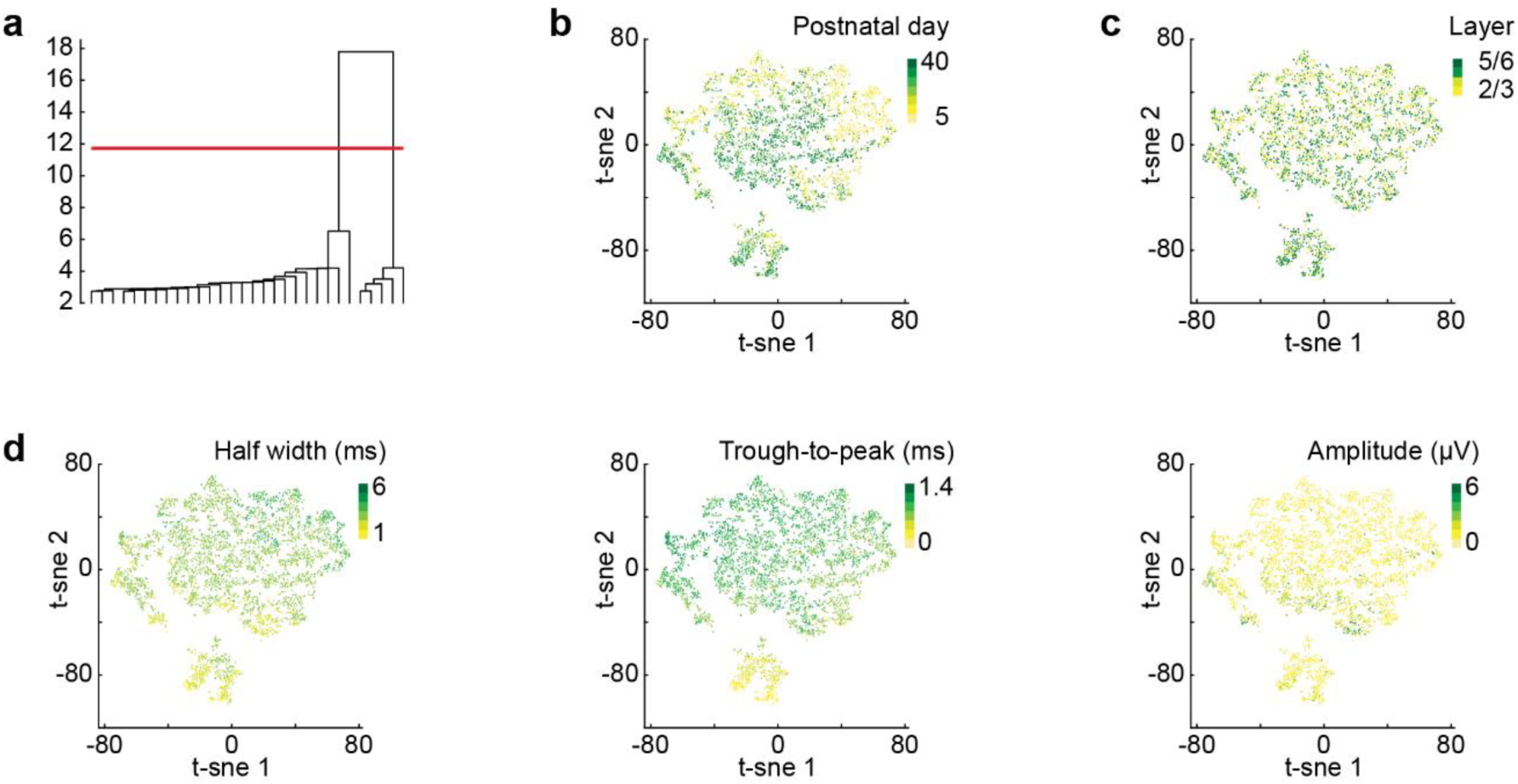
t-sne dimensionality reduction. (**a**) Dendrogram of binary hierarchical clustering after t-sne dimensionality reduction. The height of the lines represents the distance between subclusters. (**b**) Scatter plot showing the first two components of a t-sne dimensionality reduction on the mean waveforms for all units recorded from P5-40 color coded by age. (**c**) Same as (b) color coded by mPFC layer. (**d**) Same as (b) color coded by values of classic spike shape features (half width, trough to peak time, amplitude).

### Activation of L2/3 pyramidal neurons drives fast oscillations with similar acceleration across development as spontaneous activity

Besides FS interneurons, L2/3 PYRs in mPFC have been found to induce fast oscillations in the mPFC of P8-10 mice. Their non-rhythmic activation (but not activation of L5/6PYRs) drives oscillatory activity peaking within 15-20 Hz range^14,15^, similar to the peak frequency of spontaneous network activity at this age. To test if L2/3 PYRs-driven activity also accelerates with age, we optogenetically manipulated these neurons in P5-40 mice. Stable expression of the light-sensitive channelrhodopsin 2 derivate E123T T159C (ChR2(ET/TC)) restricted to about 25% of PYR in L2/3 of the mPFC was achieved by in utero electroporation (IUE) at embryonic day (E) 15.5 (Figure 3a). Optogenetic stimulation with ramps of steadily increasing light power (473 nm, 3 s) were performed during extracellular recordings in the mPFC. As previously shown, this type of stimulation activates the network without forcing a specific rhythm^14^.

**Figure 3.**
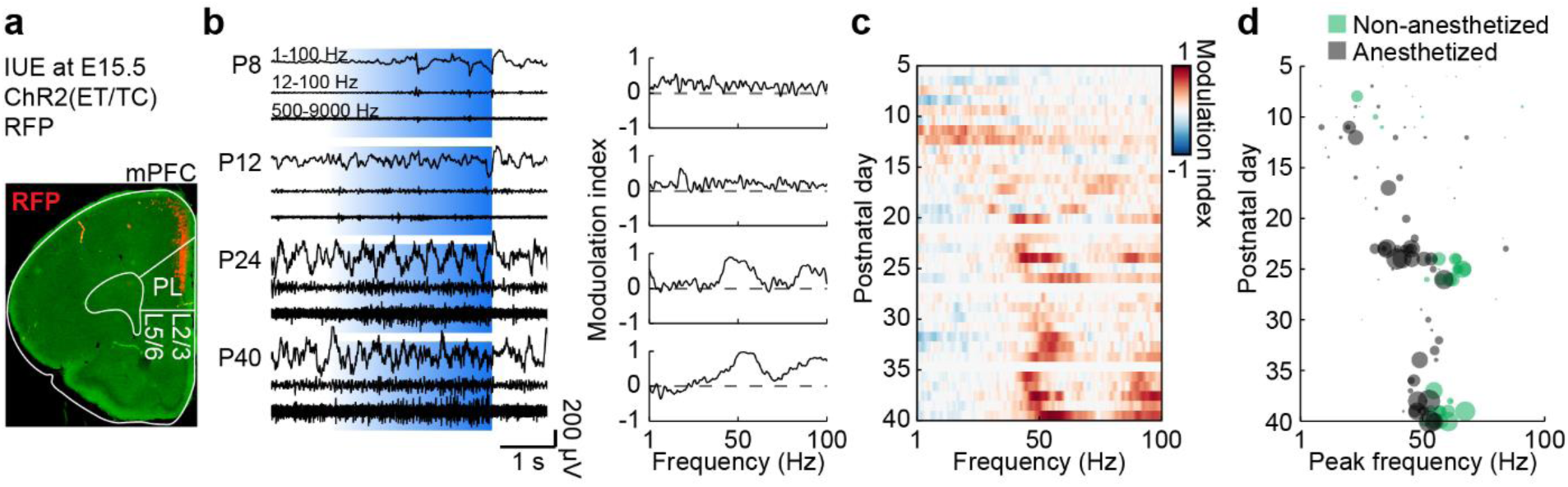
Development of L2/3 PYR-driven gamma in the mPFC. (**a**) ChR2(ET/TC)- 2A-RFP-expression in L2/3 PYRs in mPFC after IUE at E15.5 in a coronal slice of a P10 mouse. (**b**) Characteristic extracellular recordings of LFP and MUA during ramp light stimulations (473 nm, 3 s) of prefrontal L2/3 PYRs at different ages (left) and the corresponding MI of power spectra (right). (**c**) Color-coded average MI of power spectra for P5-40 mice. (**d**) Scatter plot displaying stimulus induced peak frequencies during postnatal development for anesthetized (gray, n=80) and non-anesthetized mice (green, n=20, 35 recordings). Marker size displays peak strength. (Average data is presented as median ± MAD. See supplementary table 1 for statistics.)

Similar to spontaneous activity, activating L2/3 PYR induced oscillatory activity with a gradually increasing frequency during development (Figure 3b-d). Consistent peaks in the modulation index (MI) of power spectra were detected at 15-20 Hz at the beginning of the second postnatal week and increased in frequency (Mann-Kendall trend test, p=7.69*10^−6^, n=115 recordings, tau-b 0.288) and amplitude (Mann-Kendall trend test, p=1.04*10^−9^, n=115 recordings, tau-b 0.392) until reaching stable values within 50-60 Hz at P25. Control stimulations with light that does not activate ChR2(ET/TC) (594 nm, 3 s) did not induce activity and led to the detection of unspecific peak frequencies (Mann-Kendall trend test, p=0.09, n=111 recordings, tau-b 0.111) and amplitudes (Mann-Kendall trend test, p=0.74, n=111 recordings, tau-b 0.022) (Figure 3 – figure supplement 1). Thus, L2/3 PYR driven activity in the mPFC follows the same developmental dynamics as spontaneous activity indicating the importance of L2/3 PYRs for gamma maturation.

**Figure 3 – figure supplement 1.**
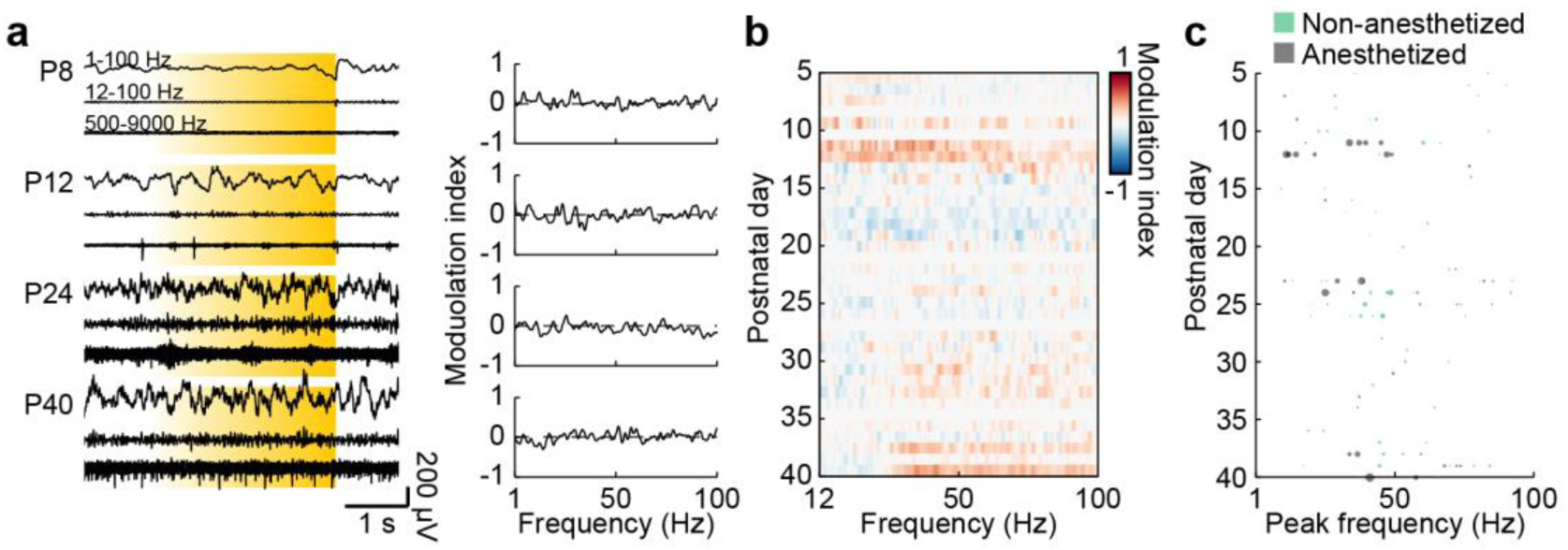
Control stimulations of L2/3 PYRs in the mPFC. (**a**) Characteristic extracellular recordings of LFP and MUA during control ramp light stimulations (594 nm, 3 s) of mPFC L2/3 PYR at different ages (left) and the corresponding MI of power spectra (right). (**c**) Color-coded average MI of power spectra for P5-40 mice. (**d**) Scatter plot displaying stimulus induced peak frequencies across age for anesthetized (gray, n=80) and non-anesthetized mice (green, n=20, 35 recordings). Marker size displays peak strength. (Average data is displayed as median ± MAD. See supplementary table 1 for statistics.)

### The rhythmicity of pyramidal cell and interneuron firing follows similar development as accelerating gamma activity in mPFC

To assess the contribution of distinct cell types to the emergence of gamma during postnatal development, we compared the firing of RS units, mainly corresponding to PYRs, and FS units, corresponding to PV-expressing interneurons, during ramp light stimulations of L2/3 PYRs in P5-40 mice.

The average firing rate of RS and FS units in the mPFC increased in response to ramp stimulation (Figure 4a). While ramp-induced firing rate changes of RS units (Mann-Kendall trend test, p=0.07, n=7 age groups, tau-b 0.619) became more prominent at older age, the firing rate changes were stable for FS units (Mann-Kendall trend test, p=0.88, n=7 age groups, tau-b 0.047) (Figure 4b). At the level of individual units, most RS units increased their firing in response to stimulation at P5-10, whereas at older age the number of activated and inactivated RS units got more balanced (Mann-Kendall trend test, p=1.52*10^−14^, n=1821 units, tau-b -0.123) (Figure 4c). In contrast, individual FS units showed a balanced distribution of activation and inactivation throughout development (Mann-Kendall trend test, p=0.91, n=225 units, tau-b -0.005), yet the low number of FS units at young age precluded clear conclusions. Thus, during early postnatal development most RS units are activated by ramp light stimulations but only moderately increase their firing rate. During late postnatal development some RS units strongly increase their firing rate, whereas others reduce their firing rate.

**Figure 4.**
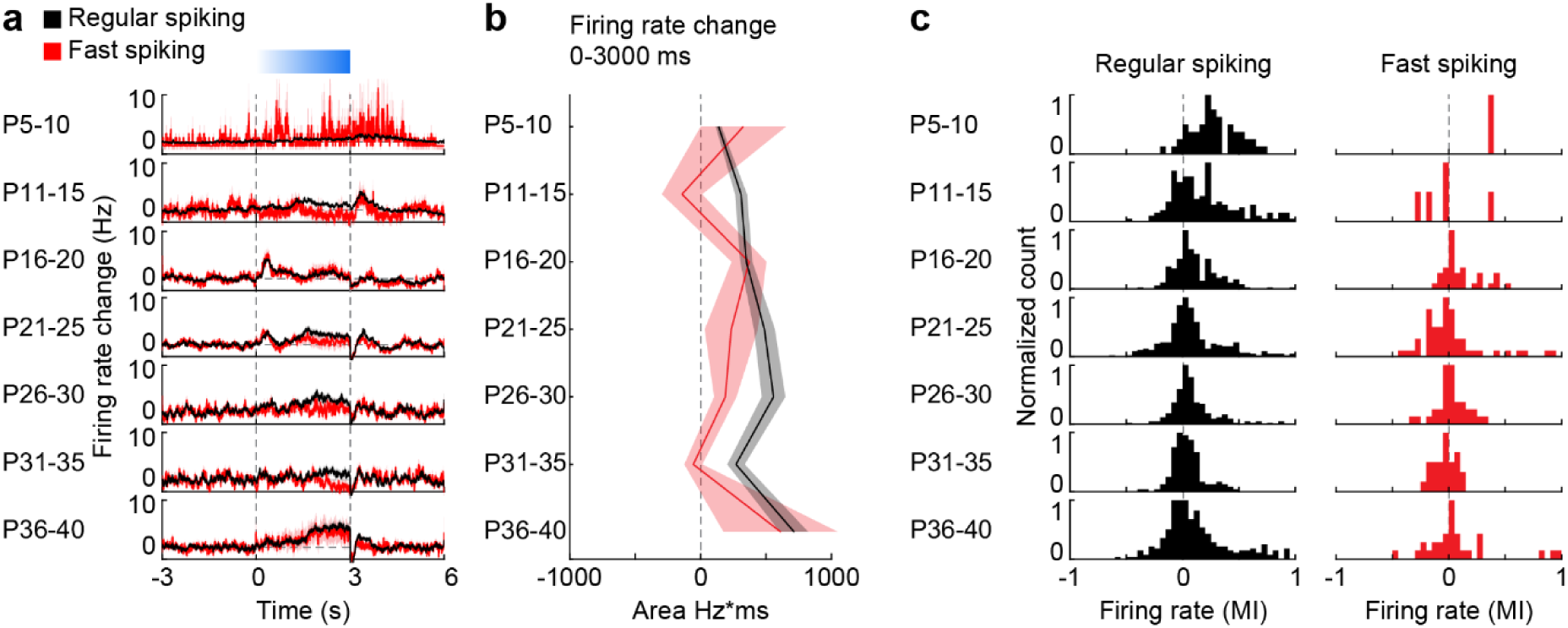
Development of RS and FS unit activity during L2/3 PYR-driven gamma in the mPFC. (**a**) Average firing rate change of RS (black) and FS (red) units in response to ramp light stimulation (3 s, 473 nm) of prefrontal L2/3 PYRs over time for different age groups. (**b**) Line plot displaying the average firing rate changes of RS and FS units during ramp light stimulation for different age groups. (**c**) Histograms of the MI of firing rates in response to ramp light stimulation for RS and FS units. (Average data is displayed as mean ± sem. See supplementary table 1 for statistics.)

Next, we tested whether RS and FS units engage in rhythmic activity and calculated autocorrelations of individual units. While no clear rhythmicity was found during spontaneous activity before stimulations (Figure 5 – figure supplement 1), autocorrelations showed that a subset of RS and FS units fire rhythmically in response to ramp light stimulation of prefrontal L2/3 PYRs (Figure 5a). The power of autocorrelations revealed that prominent rhythmic firing starts at about P15 and increases in frequency before it stabilizes at about P25 (Figure 5b). The dynamics is similar to the spontaneous and stimulated gamma activity development, indicating close interactions between RS and FS units during fast oscillations.

**Figure 5.**
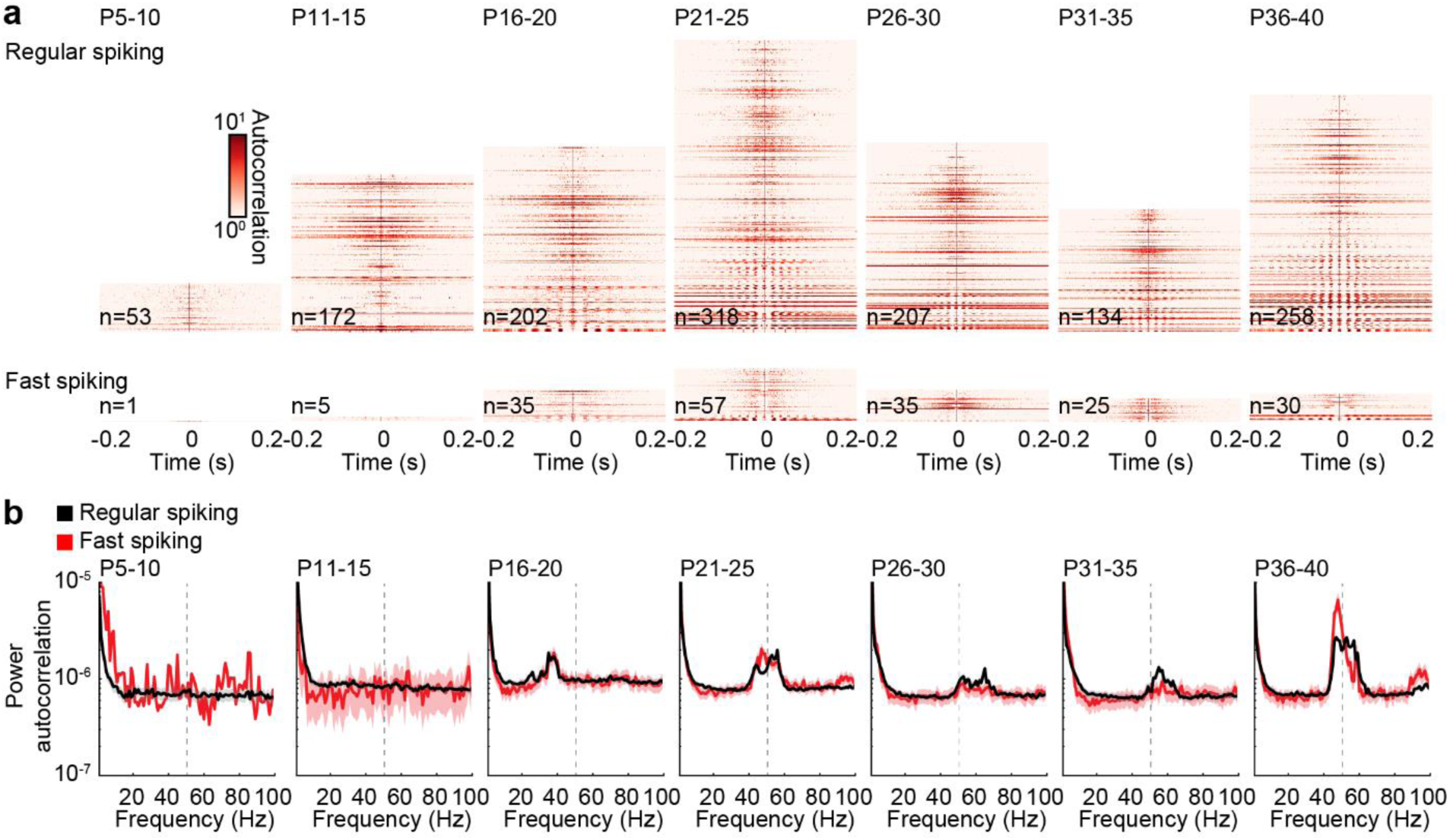
Autocorrelation of RS and FS units across age. (**a**) Color-coded autocorrelations of prefrontal RS (top) and FS (bottom) units during ramp light stimulation (3 s, 473 nm) for different age groups. Each row represents one unit. (**b**) Average autocorrelation power of RS (black) and FS (red) units during ramp light stimulation for different age groups. (Average data is displayed as mean ± sem.)

**Figure 5 – figure supplement 1.**
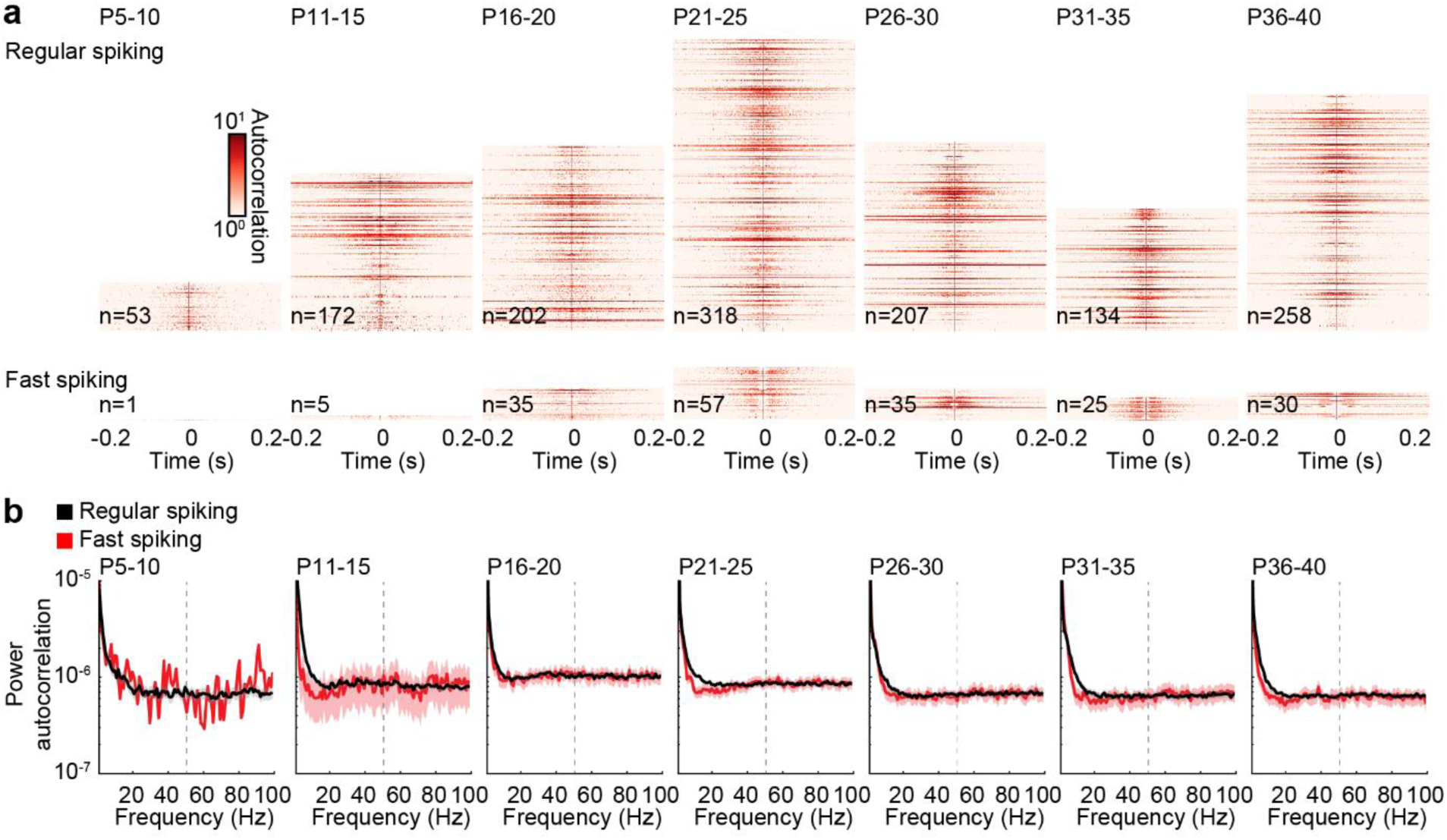
Autocorrelation of RS and FS units during spontaneous activity. (**a**) Color-coded autocorrelations of prefrontal RS (top) and FS (bottom) units for different age groups. Each row represents one unit. (**b**) Average autocorrelation power of RS (black) and FS (red) units for different age groups. (Average data is displayed as mean ± sem.)

### Inhibitory feedback maturation resembles the dynamics of gamma development

Stimulation of prefrontal L2/3 PYRs with short light pulses (3 ms, 473 nm) at different frequencies were used to test the maximal firing rates of RS and FS units in P5-40 mice. Pulse stimulations induced a short increase of firing for both RS and FS units (Figure 6a). Confirming previous results^14^, RS units did not follow high stimulation frequencies in mice younger than P10 but showed strong attenuation in the response to repetitive pulses. With ongoing development, this attenuation at high stimulation frequencies became less prominent for RS and FS units (Figure 6b). Inter-spike intervals of individual units revealed several peaks at fractions of the stimulation frequency for RS and FS, especially at higher stimulation frequencies (Figure 6 – figure supplement 1). These data suggest that individual units do not fire in response to every light pulse in the pulse train but skip some pulses.

**Figure 6.**
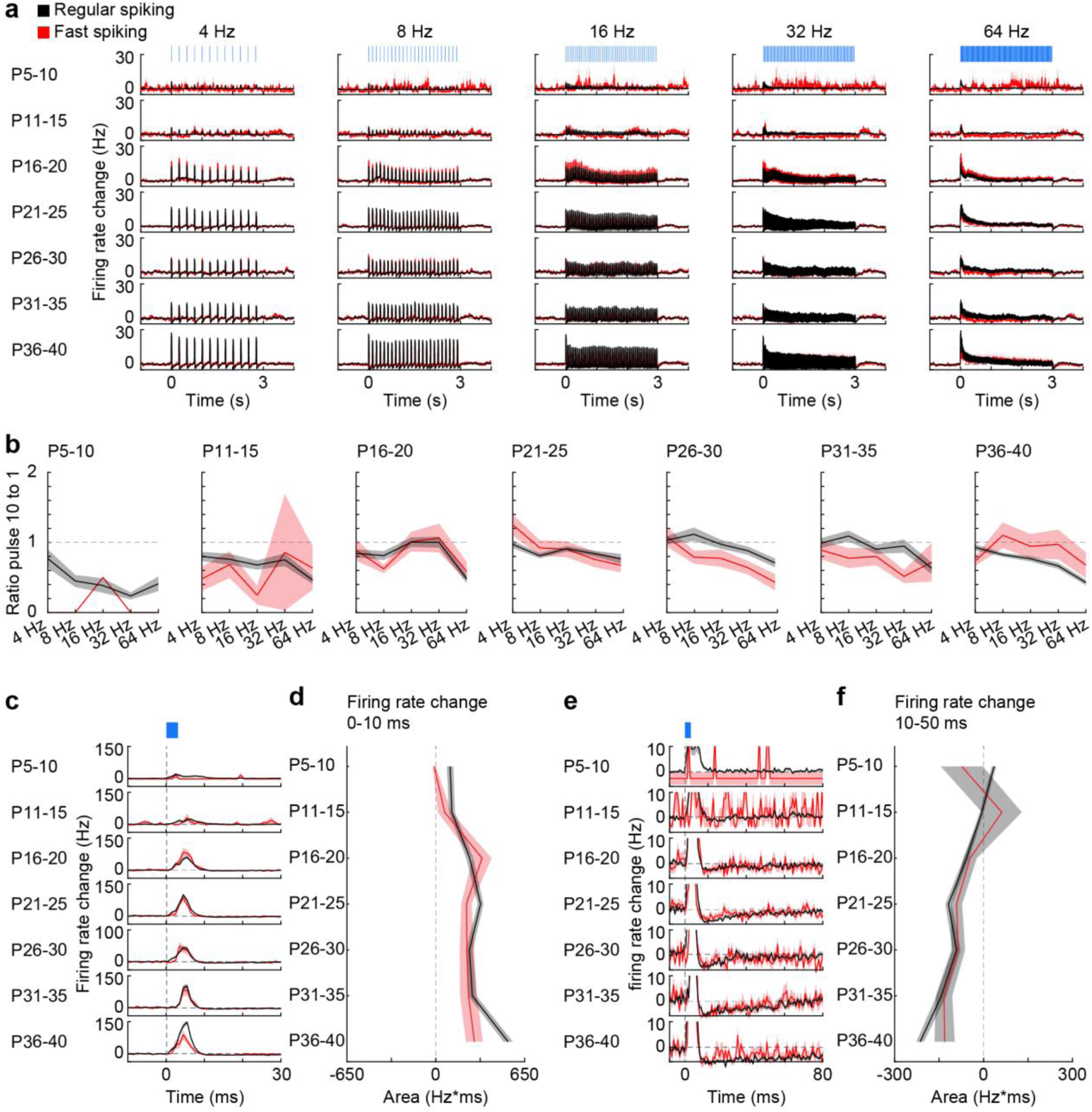
Firing of RS and FS units in response to pulse light stimulation. (**a**) Firing rate changes of prefrontal RS (black) and FS (red) units in response to repetitive pulse light stimulation (3 ms, 473 nm) of 4, 8, 16, 32, and 64 Hz averaged for different age groups. (**b**) Line plots displaying the ratio of firing rate change in response to the 10^th^ versus the 1^st^ pulse for different frequencies and age groups. (**c**) Firing rate changes of RS and FS units in response to pulse light stimulation (3 ms, 473 nm) of L2/3 PYRs averaged for different age groups. (**d**) Line plot displaying the average firing rate change of RS and FS units 0-10 ms after pulse light stimulation for different age groups. (**e**) Same as (a) displayed at longer time scale. (**f**) Same as (b) for 10-50 ms after pulse start. (Average data is displayed as mean ± sem. See supplementary table 1 for statistics.)

To assess the development of inhibitory feedback, we examined the firing rates of RS and FS units from P5-40 mice in response to individual 3 ms-long light pulses. The firing rate of RS and FS units transiently increased after pulse stimulation (Fig. 6c). This effect increased with age for RS units (Mann-Kendall trend test, p=0.04, n=7 age groups, tau-b 0.714), but not for FS units (Mann-Kendall trend test, p=0.07, n=7 age groups, tau-b 0.619) (Figure 6d). Next, we analyzed the delays of light-induced firing peaks for the two populations of units. The similar delays observed for RS and FS units suggest that the majority of RS units are non-transfected neurons, that are indirectly activated. The initial peak of increased firing was followed by reduced firing rates for RS and FS units only during late postnatal development (Figure 6e, f). The magnitude and duration of this firing depression gradually augmented with age and reached significance for RS units (Mann-Kendall trend test, RS, p=6.9*10^−3^, n=7 age groups, tau-b -0.905; FS, p=0.07, n=7 age groups, tau-b -0.619). Thus, the maturation of inhibitory feedback in response to L2/3 PYR firing in the mPFC resembles the dynamics of gamma development.

**Figure 6 – figure supplement 1.**
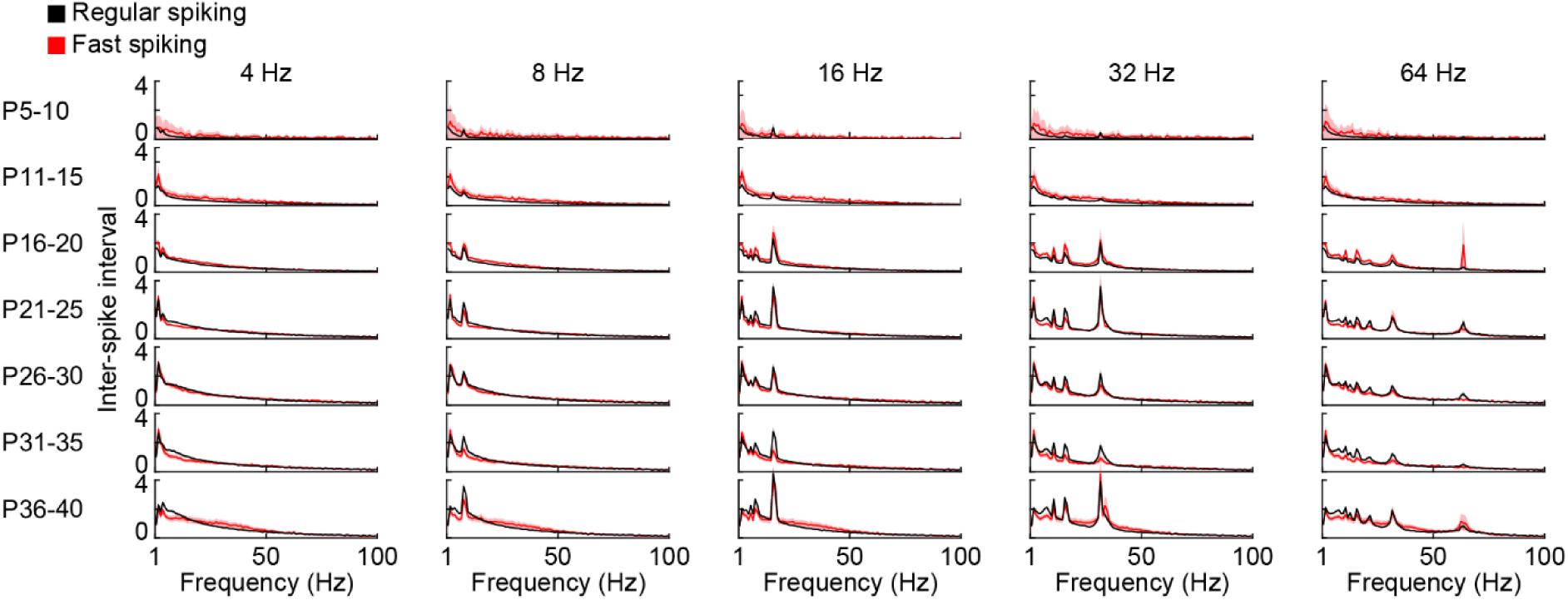
Inter-spike intervals of RS and FS units during pulse light stimulation. Inter-spike intervals of prefrontal RS (black) and FS (red) units in response to repetitive ramp light stimulation of L2/3 PYRs (3 ms, 473 nm) of 4, 8, 16, 32, and 64 Hz averaged for different age groups. (Average data is displayed as mean ± sem.)

## Discussion

Gamma oscillations result from a fine-tuned interplay between excitation and inhibition in the adult brain^16^. The mechanisms controlling the emergence of gamma activity during development are still poorly understood. Here, we reveal that the fast oscillatory activity in the mouse mPFC emerges during the second postnatal week and increases in frequency and amplitude before it stabilizes in gamma frequency range (30-80 Hz) during the fourth postnatal week. Further, we show that the functional maturation of FS PV-expressing interneurons follows similar developmental dynamics as the accelerating gamma activity. While activation of L2/3 PYR drives fast oscillatory activity throughout development, the acceleration towards higher frequencies relates to the maturation of inhibitory feedback and fast-spiking interneurons. These results demonstrate that the interplay between excitatory and inhibitory neurons is not only critical for the generation of adult gamma activity but also for its emergence during postnatal development.

Starting with the first electroencephalographic recordings, adult brain rhythms have been defined according to their frequencies and related to a specific state or task^17^. These “classical” frequency bands (i.e. delta, theta, alpha, beta, gamma) are largely preserved between different mammalian species^17,18^. However, how they emerge during development is still largely unknown. Synchronization of cortical areas in fast oscillatory rhythms starts during the first postnatal week^19^. These low-amplitude patterns are detected in the rodent mPFC as early as P5,1-2 days later compared to primary sensory cortices (S1, V1)^20–24^. However, this neonatal fast activity has a relative low frequency (around 20 Hz)^19,24^. It is organized in infrequent short bursts and its detection is hampered by the lack of a clear peak in LFP power spectra. We previously showed that in the developing mPFC the detection of prominent fast oscillations with frequencies above 12 Hz coincides with the switch from discontinuous to continuous activity^19^. These oscillations are initially within15-20 Hz frequency range that was classically defined as beta range. The present results indicate that these rhythms progressively increase their frequency and amplitude with age until they stabilize in gamma frequency range at 50-60 Hz during the fourth postnatal week. Therefore, identification of oscillatory patterns in developing circuits according to “classical” frequency bands established for adults should be avoided.

Adult gamma activity in the cerebral cortex relies on FS PV-expressing interneurons^4^. To test whether this mechanism underlies the fast rhythms in the developing brain, we developed an unbiased approach to detect FS units corresponding to putatively PV-expressing interneurons. For this, clustering of prefrontal neurons from mice of all investigated ages was performed based on a dimensionality reduction of their mean waveforms and not on pre-defined waveform features. To validate this approach, we compared the results to pre-defined waveform features typically used to identify FS units and found that they largely agree for adult mice. We demonstrate that FS units are detected in the mPFC during the second postnatal week and progressively mature until the fourth postnatal week, consistent with PV interneuron maturation^25^. The similar dynamics of FS interneuron maturation and acceleration of fast oscillatory activity supports the hypothesis that FS interneurons are key elements for prefrontal gamma development. In the absence of FS interneurons at early age, inhibitory feedback from SOM neurons – important for slow gamma activity in the adult cortex^6^ – might contribute to early oscillatory activity at frequencies within 12-20 Hz range.

While we only found minor age-dependent changes in the extracellular waveforms of RS units, an in-depth investigation of prefrontal PYRs during development identified prominent changes in their dendritic arborization, passive and active membrane properties, as well as excitatory and inhibitory inputs^26^. These changes, even though not detected with extracellular recordings, most likely contribute to the maturation of pyramidal-interneuronal interactions and finally, of gamma activity. Indeed, we found that the maturation of inhibitory feedback in response to prefrontal L2/3 PYRs stimulation follows the same dynamics as gamma development.

GABAergic transmission in the rodent cortex matures during postnatal development, reaching an adult-like state towards the end of the fourth postnatal week^27–29^. Shortly after birth, GABA acts depolarizing due to high intracellular chloride in immature neurons expressing low levels of the chloride cotransporter KCC2 relative to NKCC1^30,31^. However, this depolarization is not sufficient to trigger action potential firing and results in shunting inhibition^32^. The switch of GABA action from depolarizing to hyperpolarizing has been reported to occur during the second postnatal week^31^, coinciding with the emergence of gamma band oscillations. Moreover, the composition of GABAA-receptor subunits changes during postnatal development, causing a progressive decrease of decay-time constants of inhibitory postsynaptic currents (IPSCs) until they reach adult-like kinetics in the fourth postnatal week^25,33,34^. Simulations of neuronal networks proposed that increasing IPSCs kinetics in FS interneurons results in increasing gamma frequency^35^. The gradual increase of prefrontal gamma frequency from the second to the fourth postnatal week provides experimental evidence for this hypothesis.

In-depth understanding of the dynamics and mechanisms of gamma activity in the developing cortex appears relevant for neurodevelopmental disorders, such as schizophrenia and autisms. Both, in patients and disease mouse models, gamma oscillations have been reported to be altered, likely to reflect abnormal pyramidal-interneuronal interactions^7–9^. These dysfunction seems to emerge already during development^10–12^. Elucidating the developmental dynamics of cortical gamma activity might uncover the timeline of disease-related deficits.

## Materials and methods

### Animals

All experiments were performed in compliance with the German laws and the guidelines of the European Community for the use of animals in research and were approved by the local ethical committee (G132/12, G17/015, N18/015). Timed-pregnant mice from the animal facility of the University Medical Center Hamburg-Eppendorf were housed individually at a 12 h light/12 h dark cycle and were given access to water and food ad libitum. The day of vaginal plug detection was considered embryonic day (E) 0.5, the day of birth was considered postnatal day (P) 0. Experiments were carried out on C57Bl/6J mice of both sexes.

### In utero electroporation (IUE)

Pregnant mice (C57Bl6/J or Gad2-IRES-Cre (The Jackson Laboratory, ME, USA) crossed with Ai40 reporter line (Ai40(RCL-ArchT/EGFP)-D, The Jackson Laboratory, ME, USA)) received additional wet food daily, supplemented with 2-4 drops Metacam (0.5 mg/ml, Boehringer-Ingelheim, Germany) one day before until two days after *in utero* electroporation. At E15.5, pregnant mice were injected subcutaneously with buprenorphine (0.05 mg/kg body weight) 30 min before surgery. Surgery was performed under isoflurane anesthesia (induction 5%, maintenance 3.5%) on a heating blanket. Eyes were covered with eye ointment and pain reflexes and breathing were monitored to assess anesthesia depth. Uterine horns were exposed and moistened with warm sterile PBS. 0.75-1.25 µl of opsin- and fluorophore-encoding plasmid (pAAV-CAG-ChR2(E123T/T159C)-2A-tDimer2, 1.25 µg/µl) purified with NucleoBond (Macherey-Nagel, Germany) in sterile PBS with 0.1% fast green dye was injected in the right lateral ventricle of each embryo using pulled borosilicate glass capillaries. Electroporation tweezer paddles of 5 mm diameter were oriented at a rough 20° leftward angle from the midline of the head and a rough 10° downward angle from the anterior to posterior axis to transfect precursor cells of medial prefrontal layer 2/3 PYRs neurons with 5 electroporation pulses (35 V, 50 ms, 950 ms interval, CU21EX, BEX, Japan). Uterine horns were placed back into the abdominal cavity. Abdominal cavity was filled with warm sterile PBS and abdominal muscles and skin were sutured with absorbable and non-absorbable suture thread, respectively. After recovery from anesthesia, mice were returned to their home cage, placed half on a heating blanket for two days after surgery. Fluorophore expression was assessed at P2 in the pups with a portable fluorescence flashlight (Nightsea, MA, USA) through the intact skin and skull and confirmed in brain slices postmortem.

### Electrophysiology

#### Acute recordings

Multi-site extracellular recordings were performed unilaterally or bilaterally in the mPFC of non-anesthetized or anesthetized P5-40 mice. Mice were on a heating blanket during the entire procedure. Under isoflurane anesthesia (induction: 5%; maintenance: 2.5%), a craniotomy was performed above the mPFC (0.5 mm anterior to bregma, 0.1-0.5 mm lateral to the midline). Pups were head-fixed into a stereotaxic apparatus using two plastic bars mounted on the nasal and occipital bones with dental cement. Multi-site electrodes (NeuroNexus, MI, USA) were inserted into the mPFC (four-shank, A4×4 recording sites, 100 µm spacing, 125 µm shank distance, 1.8-2.0 mm deep). A silver wire was inserted into the cerebellum and served as ground and reference. Pups were allowed to recover for 30 min prior to recordings. For anesthetized recordings, urethane (1 mg/g body weight) was injected intraperitoneally prior to the surgery.

#### Chronic recordings

Multisite extracellular recordings were performed unilaterally in the mPFC of P23-25 and P38-40 mice. The adapter for head fixation was implanted at least days before recordings. Under isoflurane anesthesia (5% induction, 2.5% maintenance), a metal head-post (Luigs and Neumann, Germany) was attached to the skull with dental cement and a craniotomy was performed above the mPFC (0.5-2.0 mm anterior to bregma, 0.1-0.5 mm right to the midline) and protected by a customized synthetic window. A silver wire was implanted between skull and brain tissue above the cerebellum and served as ground and reference. 0.5% bupivacaine / 1% lidocaine was locally applied to cutting edges. After recovery from anesthesia, mice were returned to their home cage. After recovery from the surgery, mice were accustomed to head-fixation and trained to run on a custom-made spinning disc. For recordings, craniotomies were uncovered and multi-site electrodes (NeuroNexus, MI, USA) were inserted into the mPFC (one-shank, A1×16 recording sites, 100 µm spacing, 2.0 mm deep).

Extracellular signals were band-pass filtered (0.1-9000 Hz) and digitized (32 kHz) with a multichannel extracellular amplifier (Digital Lynx SX; Neuralynx, Bozeman, MO, USA). Electrode position was confirmed in brain slices postmortem.

### Optogenetic stimulation

Ramp (linearly increasing light power) and pulsed (short pulses of 3 ms) light stimulation was performed using an Arduino uno (Arduino, Italy) controlled laser system (473 nm / 594 nm wavelength, Omicron, Austria) coupled to a 50 µm (4 shank electrodes) or 105 µm (1 shank electrodes) diameter light fiber (Thorlabs, NJ, USA) glued to the multisite electrodes, ending 200 µm above the top recording site. Responses to stimulations were averaged for 30 repetitions.

### Histology

Mice (P5-40) were anesthetized with 10% ketamine (aniMedica, Germanry) / 2% xylazine (WDT, Germany) in 0.9% NaCl (10 µg/g body weight, intraperitoneal) and transcardially perfused with 4% paraformaldehyde (Histofix, Carl Roth, Germany). Brains were removed and postfixed in 4% paraformaldehyde for 24 h. Brains were sectioned coronally with a vibratom at 50 µm for immunohistochemistry.

#### Immunohistochemistry

Free-floating slices were permeabilized and blocked with PBS containing 0.8% Triton X-100 (Sigma-Aldrich, MO, USA), 5% normal bovine serum (Jackson Immuno Research, PA, USA) and 0.05% sodium azide. Slices were incubated over night with primary antibody rabbit-anti-parvalbumin (1:500, #ab11427, Abcam, UK) or rabbit-anti-somatostatin (1:250, #sc13099, Santa Cruz, CA, USA), followed by 2 h incubation with secondary antibody goat-anti-rabbit Alexa Fluor 488 (1:500, #A11008, Invitrogen-Thermo Fisher, MA, USA). Sections were transferred to glass slides and covered with Fluoromount (Sigma-Aldrich, MO, USA).

#### Cell quantification

Images of immunofluorescence in the right mPFC were acquired with a confocal microscope (DM IRBE, Leica, Germany) using a 10x objective (numerical aperture 0.3). Immunopositive cells were automatically quantified with custom-written algorithms in ImageJ environment. The region of interest (ROI) was manually defined over L2/3 of the mPFC. Image contrast was enhanced before applying a median filter. Local background was subtracted to reduce background noise and images were binarized and segmented using the watershed function. Counting was done after detecting the neurons with the extended maxima function of the MorphoLibJ plugin.

### Data analysis

In vivo data were analyzed with custom-written algorithms in Matlab environment. Data were band-pass filtered (500-9000 Hz for spike analysis or 1-100 Hz for local field potentials (LFP)) using a third-order Butterworth filter forward and backward to preserve phase information before down-sampling to analyze LFP.

#### Power spectral density

For power spectral density analysis, 2 s-long windows of LFP signal were concatenated and the power was calculated using Welch’s method with non-overlapping windows. Spectra were multiplied with squared frequency.

#### Modulation index

For optogenetic stimulations, modulation index was calculated as (value stimulation - value pre stimulation) / (value stimulation + value pre stimulation).

#### Peak frequency and strength

Peak frequency and peak strength were calculated for the most prominent peak in the spectrum defined by the product of peak amplitude, peak half width and peak prominence.

#### Single unit analysis

Spikes were detected and sorted with Kilosort2 in Matlab. t-sne dimensionality reduction was applied on mean waveforms of all units. Hierarchical clustering was performed to identify fast-spiking (FS) and regular-spiking (RS) units for all ages simultaneously. Auto and cross correlations were calculated before and during optogenetic ramp stimulation. Power spectral densities of mean autocorrelation were calculated per unit.

#### Statistics

Data were tested for consistent trends across age with the non-parametric Mann-Kendall trend test. Mann-Kendall coefficient tau-b adjusting for ties is reported. See supplementary table 1 for detailed statistics.

## Acknowkedgments

We thank M. Chini for helpful discussions and comments on the manuscript as well as A. Marquardt, P. Putthoff, A. Dahlmann, and K. Titze for excellent technical assistance. This work was funded by grants from the European Research Council (ERC-2015-CoG 681577 to I.L.H.-O.) and the German Research Foundation (Ha 4466/10-1, Ha4466/11-1, Ha4466/12-1, SPP 1665, SFB 936 B5 to I.L.H.-O.).

## Author contributions

S.H.B. and I.L.H.-O. designed the experiments, S.H.B. and J.A.P carried out the experiments, S.H.B. and J.A.P. analyzed the data, S.H.B., I.L.H.-O. and J.A.P interpreted the data and wrote the manuscript. All authors discussed and commented on the manuscript.

---

**Supplementary table 1.** Detailed statistical results.

|  |  | Description | Comment | Test | n | p-val | sign | taub |
| --- | --- | --- | --- | --- | --- | --- | --- | --- |
| **Figure 1** | d | Peak freq over age | per experiment - all | Mann-Kendall | 114 | 2.73E-08 | *** | 3.61E-01 |
|  | d | Peak ampl over age | per experiment - all | Mann-Kendall | 114 | 3.93E-22 | *** | 0.625472 |
|  |  | Peak freq over age | per experiment - anesthetized | Mann-Kendall | 80 | 2.30E-07 | *** | 0.404144 |
|  |  | Peak ampl over age | per experiment - anesthetized | Mann-Kendall | 80 | 1.77E-13 | *** | 0.571882 |
|  |  | Peak freq over age | per experiment - non anesthetized | Mann-Kendall | 34 | 0.02073 | * | 0.288497 |
|  |  | Peak ampl over age | per experiment - non anesthetized | Mann-Kendall | 34 | 6.13E-08 | *** | 0.671358 |
| **Figure 2** | a | PV over age | per experiment | Mann-Kendall | 38 | 1.29E-07 | *** | 0.623225 |
|  | b | SOM over age | per experiment | Mann-Kendall | 39 | 0.990283 |  | -0.00279 |
|  | h | FS proportion over age | per experiment | Mann-Kendall | 66 | 1.44E-07 | *** | 0.457857 |
|  | i | half width over age - RS | per unit | Mann-Kendall | 3172 | 1.61E-28 | *** | -0.13379 |
|  | i | half width over age - FS | per unit | Mann-Kendall | 382 | 5.17E-17 | *** | -0.29521 |
|  | i | trough to peak over age - RS | per unit | Mann-Kendall | 3172 | 4.57E-05 | *** | -0.05139 |
|  | i | trough to peak over age - FS | per unit | Mann-Kendall | 382 | 9.23E-11 | *** | -0.23647 |
|  | i | amplitude over age - RS | per unit | Mann-Kendall | 3172 | 4.45E-22 | *** | -0.11667 |
|  | i | amplitude over age<br>- FS | per unit | Mann-Kendall | 382 | 3.82E-06 | *** | -0.16274 |
| **Figure 3** | d | Peak freq over age<br>473nm | per experiment -<br>all | Mann-Kendall | 115 | 7.69E-06 | *** | 0.288624 |
|  | d | Peak MI over age<br>473nm | per experiment -<br>all | Mann-Kendall | 115 | 1.04E-09 | *** | 0.392922 |
|  |  | Peak freq over age<br>473nm | per experiment -<br>anesthetized | Mann-Kendall | 80 | 0.001207 | ** | 0.251951 |
|  |  | Peak MI over age<br>473nm | per experiment -<br>anesthetized | Mann-Kendall | 80 | 6.06E-06 | *** | 0.351405 |
|  |  | Peak freq over age<br>473nm | per experiment -<br>non anesthetized | Mann-Kendall | 35 | 0.007557 | ** | 0.33318 |
|  |  | Peak MI over age<br>473nm | per experiment -<br>non anesthetized | Mann-Kendall | 35 | 0.000126 | *** | 0.475853 |
| **Figure 3 -**<br>**supplement 1** | c | Peak freq over age<br>594nm | per experiment -<br>all | Mann-Kendall | 111 | 0.092354 |  | 0.110678 |
|  | c | Peak MI over age<br>594nm | per experiment -<br>all | Mann-Kendall | 111 | 0.741814 |  | 0.02177 |
|  |  | Peak freq over age<br>594nm | per experiment -<br>anesthetized | Mann-Kendall | 76 | 0.096874 |  | 0.133006 |
|  |  | Peak MI over age<br>594nm | per experiment -<br>anesthetized | Mann-Kendall | 76 | 0.811538 |  | 0.019391 |
|  |  | Peak freq over age<br>594nm | per experiment -<br>non anesthetized | Mann-Kendall | 35 | 0.52095 |  | 0.081591 |
|  |  | Peak MI over age<br>594nm | per experiment -<br>non anesthetized | Mann-Kendall | 35 | 0.988098 |  | -0.00369 |
| **Figure 4** | b | RS rate change<br>over age | per age group | Mann-Kendall | 7 | 0.071505 |  | 0.619048 |
|  | b | FS rate change<br>over age | per age group | Mann-Kendall | 7 | 0.880616 |  | 0.047619 |
|  | c | RS MI hist over age | per unit | Mann-Kendall | 1821 | 1.52E-14 | *** | -0.12261 |
|  | c | FS MI hist over age | per unit | Mann-Kendall | 225 | 0.906635 |  | 0.005438 |
| **Figure 6** | d | RS rate change over age | per age group | Mann-Kendall | 7 | 0.035498 | * | 0.714286 |
|  | d | FS rate change over age | per age group | Mann-Kendall | 7 | 0.071505 |  | 0.619048 |
|  | f | RS rate change over age | per age group | Mann-Kendall | 7 | 0.006864 | ** | -0.90476 |
|  | f | FS rate change over age | per age group | Mann-Kendall | 7 | 0.071505 |  | -0.61905 |

## References

1. Singer, W. Neuronal oscillations: unavoidable and useful? Eur. J. Neurosci. 48, 2389–2398 (2018).

2. Cardin, J. A. Snapshots of the Brain in Action: Local Circuit Operations through the Lens of γ Oscillations. J. Neurosci. 36, 10496–10504 (2016).

3. Sohal, V. S. How Close Are We to Understanding What (if Anything) γ Oscillations Do in Cortical Circuits? J. Neurosci. 36, 10489–10495 (2016).

4. Cardin, J. A. et al. Driving fast-spiking cells induces gamma rhythm and controls sensory responses. Nature 459, 663–667 (2009).

5. Chen, G. et al. Distinct Inhibitory Circuits Orchestrate Cortical beta and gamma Band Oscillations. Neuron 96, 1403–1418.e6 (2017).

6. Veit, J., Hakim, R., Jadi, M. P., Sejnowski, T. J. & Adesnik, H. Cortical gamma band synchronization through somatostatin interneurons. Nat. Neurosci. 20, 951–959 (2017).

7. Cho, K. K. A. et al. Gamma Rhythms Link Prefrontal Interneuron Dysfunction with Cognitive Inflexibility in Dlx5/6+/-Mice. Neuron 85, 1332–1343 (2015).

8. Cao, W. et al. Gamma Oscillation Dysfunction in mPFC Leads to Social Deficits in Neuroligin 3 R451C Knockin Mice. Neuron 97, 1253–1260.e7 (2018).

9. Rojas, D. C. & Wilson, L. B. Gamma-band abnormalities as markers of autism spectrum disorders. Biomark. Med. 8, 353–368 (2014).

10. Chini, M. et al. Resolving and Rescuing Developmental Miswiring in a Mouse Model of Cognitive Impairment. Neuron 105, 60–74.e7 (2020).

11. Richter, M. et al. Altered TAOK2 activity causes autism-related neurodevelopmental and cognitive abnormalities through RhoA signaling. Mol. Psychiatry 24, 1329–1350 (2019).

12. Hartung, H. et al. From Shortage to Surge: A Developmental Switch in Hippocampal-Prefrontal Coupling in a Gene-Environment Model of Neuropsychiatric Disorders. Cereb. Cortex N. Y. N 1991 26, 4265–4281 (2016).

13. Chini, M. et al. Neural Correlates of Anesthesia in Newborn Mice and Humans. Front. Neural Circuits 13, (2019).

14. Bitzenhofer, S. H. et al. Layer-specific optogenetic activation of pyramidal neurons causes beta-gamma entrainment of neonatal networks. Nat. Commun. 8, 14563 (2017).

15. Bitzenhofer, S. H., Ahlbeck, J. & Hanganu-Opatz, I. L. Methodological Approach for Optogenetic Manipulation of Neonatal Neuronal Networks. Front. Cell. Neurosci. 11, 239 (2017).

16. Atallah, B. V. & Scanziani, M. Instantaneous modulation of gamma oscillation frequency by balancing excitation with inhibition. Neuron 62, 566–577 (2009).

17. Buzsáki, G. & Draguhn, A. Neuronal Oscillations in Cortical Networks. Science 304, 1926–1929 (2004).

18. Buzsáki, G., Logothetis, N. & Singer, W. Scaling Brain Size, Keeping Timing: Evolutionary Preservation of Brain Rhythms. Neuron 80, 751–764 (2013).

19. Brockmann, M. D., Pöschel, B., Cichon, N. & Hanganu-Opatz, I. L. Coupled Oscillations Mediate Directed Interactions between Prefrontal Cortex and Hippocampus of the Neonatal Rat. Neuron 71, 332–347 (2011).

20. Minlebaev, M., Colonnese, M., Tsintsadze, T., Sirota, A. & Khazipov, R. Early Gamma Oscillations Synchronize Developing Thalamus and Cortex. Science 334, 226–229 (2011).

21. Yang, J.-W. et al. Thalamic network oscillations synchronize ontogenetic columns in the newborn rat barrel cortex. Cereb. Cortex N. Y. N 1991 23, 1299–1316 (2013).

22. Dupont, E., Hanganu, I. L., Kilb, W., Hirsch, S. & Luhmann, H. J. Rapid developmental switch in the mechanisms driving early cortical columnar networks. Nature 439, 79–83 (2006).

23. Shen, J. & Colonnese, M. T. Development of Activity in the Mouse Visual Cortex. J. Neurosci. 36, 12259–12275 (2016).

24. Yang, J.-W., Hanganu-Opatz, I. L., Sun, J.-J. & Luhmann, H. J. Three Patterns of Oscillatory Activity Differentially Synchronize Developing Neocortical Networks In Vivo. J. Neurosci. 29, 9011–9025 (2009).

25. Okaty, B. W., Miller, M. N., Sugino, K., Hempel, C. M. & Nelson, S. B. Transcriptional and electrophysiological maturation of neocortical fast-spiking GABAergic interneurons. J. Neurosci. Off. J. Soc. Neurosci. 29, 7040–7052 (2009).

26. Kroon, T., van Hugte, E., van Linge, L., Mansvelder, H. D. & Meredith, R. M. Early postnatal development of pyramidal neurons across layers of the mouse medial prefrontal cortex. Sci. Rep. 9, (2019).

27. Le Magueresse, C. & Monyer, H. GABAergic Interneurons Shape the Functional Maturation of the Cortex. Neuron 77, 388–405 (2013).

28. Butt, S. J., Stacey, J. A., Teramoto, Y. & Vagnoni, C. A role for GABAergic interneuron diversity in circuit development and plasticity of the neonatal cerebral cortex. Curr. Opin. Neurobiol. 43, 149–155 (2017).

29. Lim, L., Mi, D., Llorca, A. & Marín, O. Development and functional diversification of cortical interneurons. Neuron 100, 294–313 (2018).

30. Rivera, C. et al. The K+/Cl-co-transporter KCC2 renders GABA hyperpolarizing during neuronal maturation. Nature 397, 251–255 (1999).

31. Ben-Ari, Y., Khalilov, I., Kahle, K. T. & Cherubini, E. The GABA excitatory/inhibitory shift in brain maturation and neurological disorders. Neurosci. Rev. J. Bringing Neurobiol. Neurol. Psychiatry 18, 467–486 (2012).

32. Kirmse, K., Hübner, C. A., Isbrandt, D., Witte, O. W. & Holthoff, K. GABAergic Transmission during Brain Development: Multiple Effects at Multiple Stages. The Neuroscientist 24, 36–53 (2018).

33. Bosman, L. W. J., Heinen, K., Spijker, S. & Brussaard, A. B. Mice lacking the major adult GABAA receptor subtype have normal number of synapses, but retain juvenile IPSC kinetics until adulthood. J. Neurophysiol. 94, 338–346 (2005).

34. Laurie, D. J., Wisden, W. & Seeburg, P. H. The distribution of thirteen GABAA receptor subunit mRNAs in the rat brain. III. Embryonic and postnatal development. J. Neurosci. Off. J. Soc. Neurosci. 12, 4151–4172 (1992).

35. Doischer, D. et al. Postnatal Differentiation of Basket Cells from Slow to Fast Signaling Devices. J. Neurosci. 28, 12956–12968 (2008).

